# Structural-guided fragment-based drug discovery applied to antitoxin, MAB3862 opens a new possibility of exploring the Toxin and Antitoxin for antibiotics

**DOI:** 10.1101/2022.05.14.491971

**Authors:** So Yeon Kim, Hyun-Jong Eun, JooYeon Lee, Wonsuk Lee, Sherine Elizabeth Thomas, Paul Brear, Sundeep Chaitanya Vedithi, Chris Abell, Bong-Jin Lee, Tom L. Blundell

## Abstract

*Mycobacterium abscessus* (Mab) is a rapidly growing multidrug-resistant species among nontuberculous mycobacteria (NTM). Pulmonary infections caused by *M. abscessus* are difficult to treat and often result in an accelerated condition and premature death of immunosuppressed patients such as those with cystic fibrosis, putting them at a greater risk of other infections and increasing the urgency of developing a novel class of antibiotics. Here, we explore the use of toxin and antitoxin (TA) as an interesting and promising new class of target in a fragment-based drug-discovery approach. A de novo structure of Mab3862, an antitoxin of type 2 TA class in *M. abscessus* was elucidated and followed by drug discovery work. Very small molecules (fragments) are used to bind the antitoxin and then elaborated into drug-sized molecule that potentially trigger conformational change to prevent formation of toxin-antitoxin complex. Biophysical screening methods and binding-mode guidance from *in silico* docking were conducted. Grouping of fragments based on the binding site, creating a pharmacophore model, can facilitate further studies for rational design of inhibitors. Although targeting the TA complex for developing antibiotics is relatively novel and challenging, this could possibly open the gate for exploring it as potential drug target. Targeting one of the TA pairs might not be completely bactericidal. However, using this approach strategically with other antibiotics for synergic effect might be effective for patients with a persistence phenotype requiring prolonged therapy.

## 1. Introduction

*Mycobacterium abscessus* (Mab) has been recognised as a challenging respiratory pathogen among the non-tuberculous mycobacteria (NTM), often associated with poor outcomes of therapy and a high cost incurred due to prolonged therapy [1]. Although its acquisition was thought to occur through environmental exposure (e.g. soil or water), it has become evident that indirect person-to-person transmission may also play an important role in transmitting infection among immunosuppressed patients [2]. Mab infection results in accelerated inflammatory lung damage, impaired quality of life and premature death. Thus, lung disease caused by *M. abscessus* can be considered a chronic infection that is rarely cured by currently available drugs [3].

One of the challenges of treating prolonged bacterial infection is subgroups of persister cells survive through a course of treatment and re-thrive when the antibiotic stress is lifted [4]. The persistence phenotype is an epigenetic change by a subgroup of bacterial colonies, characterized by slow growth and low metabolism, coupled with an ability to survive antibiotic treatment [5]. It is now clear that persisters are likely to be a cause of at least some of deaths since they underlie latent infections and post-treatment relapse, therefore, it appears as a major concern of public health [6]. Furthermore, repeated rounds of antibiotic treatment increase the likelihood of acquisition of resistance by bacteria to existing antibiotics.

Toxin-antitoxin (TA) systems are involved in essential pathways of bacterial life such as multidrug tolerance, biofilm formation, and arrest of cellular growth under stress conditions so triggering persistence in bacteria [7]. The TA operon consists of a toxin genes and an antitoxin gene, where the antitoxin antagonises toxic effects of the toxin by forming a complex to neutralise it [8]. There are six types of TA classes that differ in composition of antitoxin (either RNA or protein) and/or function of the toxin. Among those, type 2 is the most well understood class and is highly associated with inducing a persistence phenotype [7]. When bacteria encounter a stressful environment, antitoxin becomes unstable and is degraded by host proteases, releasing toxin to participate in post-segregational killing (PSK), abortive infection, biofilms, and persister cell formation [7–9].

Furthermore, there are no homologues of TA genes found in humans since TA is mostly spread in bacteria and some species in archaea. This means the drug against TA is likely to have a higher specificity and fewer side effects caused by off-target interactions. Also, being a novel antibiotic target implies there is no pre-existing resistance as yet developed against this type of target [10]. Overall, TAs are ideal targets which have potential in drug discovery still to be explored [8].

One of the proposed approaches for targeting the toxin-antitoxin complex is to identify peptides or small molecules that bind to antitoxin with a higher affinity to prevent TA complex formation, so that free toxin will then interact with its cellular targets for various reactions [11,12]. Here, we employ fragment-based drug discovery (FBDD) for exploring chemical space efficiently with an in-house library, a small set around 480 fragments. The criteria for selection of these fragments is driven mainly by the Rule of Three: molecular mass less than 300 Da, the number of hydrogen bond donors and acceptors each are less or equal to 3 and so on as developed in Astex Pharma [13]. Fragment-based lead discovery deals with low molecular mass and low affinity molecules, so that later in development stage they can be optimized into drug leads with optimal physicochemical features and high ligand efficiency (LE). This fragment-based approach has been very successful, leading to six FDA approved and launched drugs. Moreover, the approach has been successfully employed on what were previously considered to be challenging targets based on traditional approaches [14].

## 2. Materials & Methods

### 2.1 PCR and molecular cloning

The *Mab3862&3863* genes were amplified from *Mycobacterium abscessus* (ATCC 19977) genomic DNA using the following customised primers (Sigma):

For mab3862 (antitoxin)

Forward Primer (NcoI site) : 5’-GGAATT**CCATGG**CGACCGATGCAGATG-3’ and Reverse Primer (Hindiii site) : 5’-CGC**AAGCTT**TTAATCTTTGCTCGGACG -3’ For mab3863 (toxin)

Forward Primer (BamHi site) : 5’-GGAATT**GGATCC**ATGGCGCCTGTTACCAATGTT -3’

Reverse Primer (Hindiii site) : 5’-CGC**AAGCTT**TTAGCTGCGAACCAGTT-3’

The purified PCR products and vector (pHAT4 and pET28a) were subjected to restriction digestion with endonucleases (ThermoScientific). pHAT4 (https://www.addgene.org/112585/) was kindly offered by Hyvonen’s group [15] whereas pet28a is available from Addgene (https://www.addgene.org/vector-database/2565/).

The ligation of digested insert and vector was performed using T4 DNA ligase (New England Biolabs) by incubation at room temperature for 30 min.

Both plasmids were co-transformed into *E. coli* DH5α competent cells by the heat-shock method and plated on selective LB agar plates and incubated at 37°C. Colonies with desirable plasmids were isolated & cultured and purified to check the integrity by sequencing (DNA Sequencing Facility, Department of Biochemistry, Cambridge)

### 2.2 Expression and purification of full length of Mab3862 and Mab3863

*E. coli* BL21 (DE3) strain containing mab3862·pHAT4 and mab3863·pET28a were grown overnight at 37°C in primary culture of LB-media. This seed stage culture was used to inoculate 6 L of 2XYT media until optical density (A_600nm_) reached 0.6-0.8. The expression of recombinant construct was induced by the addition of Isopropyl β-D-1-thiogalactopyranoside (IPTG) to a final concentration of 1mM and further allowed to grow for 4 hours at 37°C. Cells were harvested by centrifugation at 4°C for 25 min at 4200 g and the pellet was re-suspended in buffer A (50 mM Tris-HCl pH 8.0, 500 mM NaCl, 0.1% w/v Sodium azide). The cells were lysed by sonication (Branson). The lysate clarified by centrifugation at 4°C for 40 min at 25,568 g, was passed through a pre-equilibriated (with buffer A), 5 ml pre-packed Nickel-sepharose column (HiTrap IMAC FF, Cytiva). The column was washed with buffer A and the bound protein was eluted using buffer B (50 mM Tris-HCl pH 8.0, 500 mM NaCl, 0.1% w/v Sodium azide and 500 mM Imidazole). Eluates from IMAC column were pooled and diluted to reduce the NaCl and imidazole level below 50mM for anion exchange (HiTrap Q Hi Performance, Cytiva) and eluted by salt gradient from 0 to 2M NaCl (HiTrap Q Hi Performance, Cytiva).

After anion exchange, eluate was concentrated using PES membrane centrifugal concentrator (Amicon) to reach an appropriate concentrated volume (<2ml) and was loaded onto a pre-equilibrated (with buffer D: 50 mM Tris-HCl pH 8.0, 150 mM NaCl) 120 ml Superdex200 16/600 column (GE Healthcare). 2 ml fractions were collected and analysed on an SDS-PAGE gel. Only those fractions corresponding to pure target proteins were pooled and concentrated to 20mg/ml for TA co-purification while purifying either toxin or antitoxin alone was 10mg/ml. Purified samples were flash frozen in liquid nitrogen and stock was stored at −80°C. Identification of protein was further aided by the analysis of liquid chromatography-mass spectrometry (LC-MS).

### 2.3 Seleno-methyl (Sel-met) protein purification of full length of Mab3862

The same construct (mab3862·pHAT4) were used to transform prototroph *E. coli* BL21 (DE3) to purify Sel-met substituted protein with repressing host methionine biosynthesis (see reference for detailed methods [16]). Primary cultures were incubated at 37ºC overnight in LB media with seed stock and the cell were harvested next day by centrifugation (4200 g for 25 mins) and transferred to new media with Seleno-Met Medium Base (MD12-501, Molecular Dimensions), 60mg of L-(+)-seleno-methionine and 100mg of threonine, lysine, phenylalanine each and 50mg of leucine, isoleucine, valine each per 1 litre of culture. The rest of culture & purification methods are same as that of native protein.

### 2.4 Thermal Shift Assays

Thermal shift assays were carried out in a 96-well format with each well containing 25 μl of reaction mixture of 10 μM protein in buffer (50 mM Tris-HCl pH 8.0, 150 mM NaCl), 5 mM compound, 5 % DMSO and 5x Sypro orange dye. The measurements were performed in a Biorad-CFX connect thermal cycler using the following program: 25°C for 10 min followed by a linear increment of 0.5°C every 30 seconds to reach a final temperature of 95°C. The results were analysed and plotted using Microsoft Excel.

### 2.5 Surface Plasmon Resonance

Biacore T200 using Series S sensor CM7 chip (Cytiva) immobilised with antitoxin, Mab3862 through amine coupling. For immobilisation, the contact time between protein and chip surface were 120 sec at the flow rate of 10μl/min in 10mM of sodium acetate pH 7.0 (adjusted with HCl) buffer at the protein concentration of 30μg/ml. The screening buffer was sodium potassium phosphate pH 6.6 (adjusted with HCl) 10mM, NaCl 150mM, Sodium azide 0.1% w/v and DMSO 1%. During the screening, contact time of each fragment (final concentration 1mM) was 60 sec at the rate of 10μl/min. At a higher concentration SPR screening, the contact time was 40 sec at the rate of 10μl/min of each fragment concentration of 3mM. For every cycle tested, the wash was performed with 3% DMSO for 30 sec at 30μl/min on flow channel while the rest of system is being washed with 50% DMSO. Solvent corrections were performed before and after each plate (every 80 fragments) for screening with DMSO ranging from 0.5-5% in a running buffer.

### 2.6 Crystallisation of Mab3862

Crystals for this experiment were grown at 25°C in 48-well sitting drop plates (Swiss CDI) in the following condition: 2.0M Ammonium sulphate, 100mM Sodium acetate pH 3.5 (adjusted with HCl). Crystals were cryo-protected using 25% ethylene glycol before flash frozen by liquid nitrogen. Drop volumes of protein and reservoir were 10μl and 5μl each (2:1 ratio).

### 2.7 Data Collection and Processing

X-ray data sets for Mab3862 were collected on i04 beamlines at the Diamond Light Source in the UK. The crystals were flash-cooled in cryo-protectant containing either 25% ethylene glycol or 30% glycerol. The data sets were collected using the rotation methods at wavelength of 0.9795 Å, Omega start: 0°, Omega Oscillation: 0.1°, total oscillation: 360°, total images: 3600, exposure time: 0.01s. The experimental intensities were processed to 2.7 AL and the diffraction images were processed using AutoPROC [17] utilizing XDS [18]for indexing, integration, followed by POINTLESS [19], AIMLESS [20]and TRUNCATE [21] programs from CCP4 Suite [22].

### 2.8 Crystal Structure Solution and refinement

The *Mycobacterium abscessus* Mab3862 antitoxin structures were solved by Autosol [23] for picking up heavy atom (Selenium in Sel-met) signals then build the model by Autobuild [24] with a primary sequence and Sel-met sites. Subsequent models were rebuilt using COOT interactive graphic program [25]. Then finally the structure was refined using Phenix refinement and validation programme [26] before deposition into PDB (https://www.rcsb.org/).

### 2.9 (Virtual screening) Protein and ligand preparation

Mab3862 structure (7R22) contained three seleno-methionine from SAD experiments. Firstly, they were all converted into methionine. Hydrogens were added where required to all residue models. Protein residues charge were adjusted at pH□=□7 to reflect physiologically relevant state using Protein Preparation Wizard in Maestro [27]. The preparation of in-house fragment library for 2D-to-3D conversion was performed and then subsequently addition of hydrogen, neutralisation of charged group at physiologically relevant pH, generation of ionisation states [28], low-energy ring conformation and tautomer followed by using the OPLS-2005 [29] were completed. After ligand preparation process, total number of molecules reached up to ∼780 because of different combinations of tautomer and enantiomer of each fragment.

### 2.10 (Virtual screening) Site mapping

SiteMap, a module of Maestro, was used to outline the protein binding sites of the complex target [30]. A grid was placed over the whole target protein by SiteMap. Vertices were allocated inside concavities and were called ‘site points’. Druggable sites were identified according to the SiteScore.

### 2.11 (Virtual screening) Ligand docking

Virtual Screening of the fragment library was carried out using the Glide’s high-throughput virtual screening (HTVS) [31] docking module for the *site 1* and *site 2* of the Mab3862 structure which was previously prepared using Protein Preparation Wizard tool. Ligands that are able to interact with these residues were ranked by their Glide Score [31].

### 2.12 (Virtual screening) Pharmacophore model generation

A visual examination from the top ranked Glide score to the bottom allows fragments that commonly interact with the overlapping site to be assembled together. Then pharmacophore hypotheses were generated using PHASE based on the e-pharmacophore [32,33].

### 2.13 Visual and analysis tools for figures

Clustal Omega for multiple alignment on sequence [34] is from EMBL-EBI website (https://www.ebi.ac.uk/Tools/msa/clustalo/). DNA B-form structure was computationally generated (http://www.scfbio-iitd.res.in/software/drugdesign/bdna.jsp). PyMol [35], Schrodinger Maestro [36] and Arpeggio [37] were used for visual presentation & inspection in this paper.

## 3. Results and discussion

### 3.1 Mab3862 structure

The asymmetric unit of the crystal contains one protomer of Mab3862, which forms a dimer related by a 2-fold symmetry. Its primitive cubic unit cell has each length corresponding to 106.89 Å (a=b=c) with space group of P4_1_32. The structure was refined with acceptable statistics; a final R-factor (0.299) and R-free (0.347). The details of data collection and refinement are presented in Table 1. The electron density for the N-terminal region, (residues from 1 to 40) corresponding mostly to an affinity tag, is not visible. Each protomer (PDB id 7R22) forms overall fold of α1-α2-α3-α4-loop1-α5 shown in Fig. 1a. It primarily consists of an alpha helix (70%), and the rest is a disordered loop connecting between helices. A Mab3862 homodimer has dimensions approximately 20Å by 40Å by 55Å (Fig. 1b), with 36% residues interacting at the interface between the two protomers. Complex Formation Significance Score (CSS) of Mab3862 protomers is computed to be 1.0 (range 0-1) [38] implying that its multimeric state has a biological significance rather than being an artifact of crystal packing. Two helices formed by α1 and α5 constitute a significant part of the dimeric interface of Mab3862. Each helix aligns antiparallel to its cognate helix of the other subunit. The physicochemical features of the dimeric interface are shown in Table 2. Additionally, an unusual feature noted between *α*1 and *α*2 is an inverse gamma turn exhibited in Leu17 (*i*), Ser18 (*i+1*) and Asp19 (*i+2*); there are hydrogen bonds between the carbonyl oxygen of Leu17 and the main chain nitrogen atom of Asp19 as well as Arg23 (Fig. 1c).

**Fig. 1.**
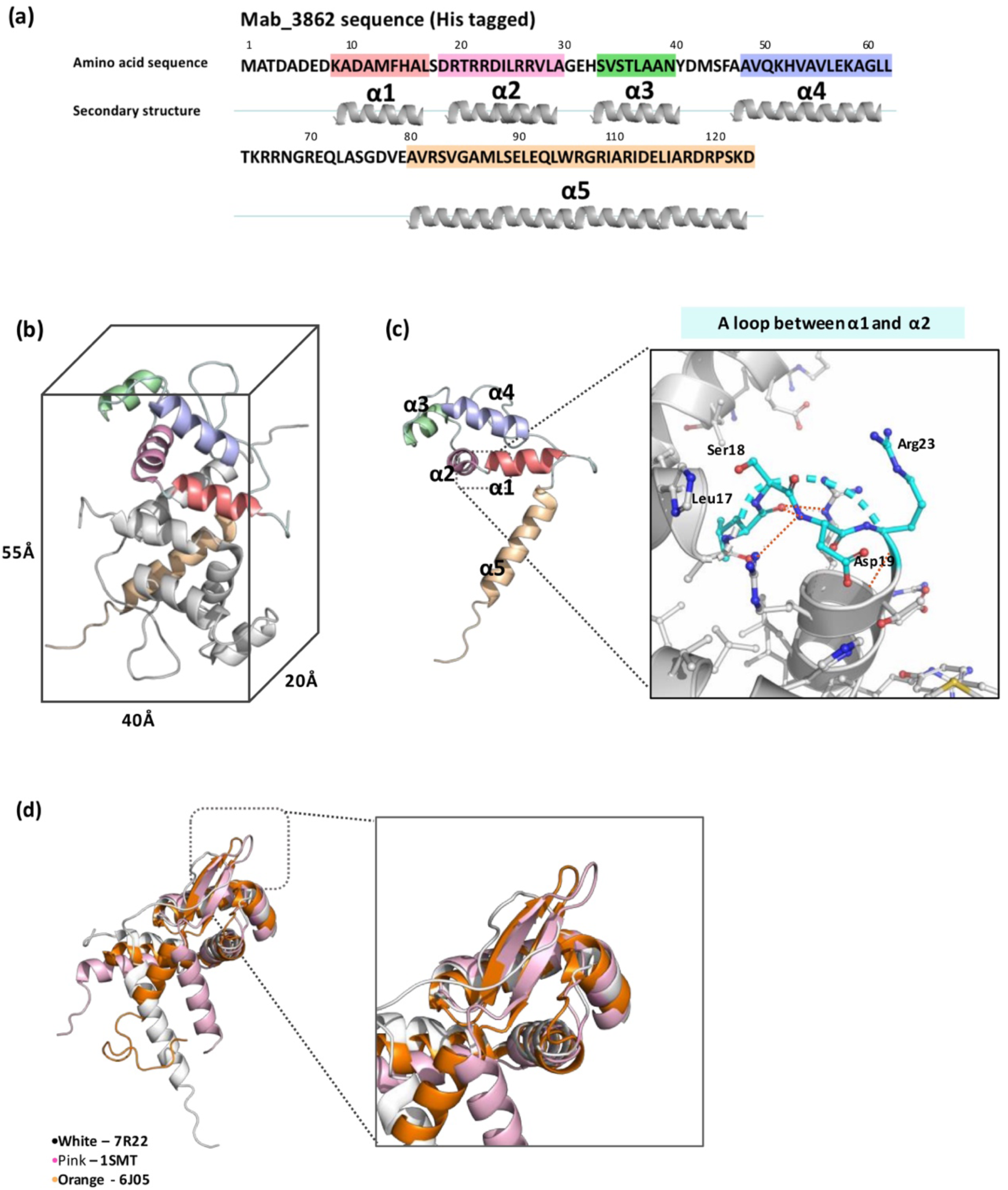
Sequence and structure of Mab3862 (PDB id 7R22). a) Sequence and secondary structure annotation of Mab3862. b) Ribbon diagram of Mab3862 dimer with secondary structural units (α1-α2-α3-α4-loop-α5) and approximate dimensions. c) An annotated Mab3862 protomer focusing on the inverse gamma turn highlighted in cyan. The hydrogen bonds are marked with orange dashed lines. (d) Comparison of the structures of Mab3862 (*Mycobacterium abscessus*, white), SmtB (*Synechococcus elongatus*, pink) and ArsR (*Acidithiobacillus ferrooxidans*, orange). The general folds are similar with the exception of the regions highlighted in the box; SmtB and ArsR form antiparallel beta strands whereas Mab3862 does not have a defined secondary structure in this region.

### 3.2 ArsR/SmtB protein family

Sequence- (pBLAST) and structure- based (DALI server) search suggest that Mab3862 belongs to the ArsR/SmtB family with a typical motif of wHTH (winged helix-turn-helix) [39,40]. The ArsR/SmtB family of metalloregulatory transcriptional regulators represses the expression of operons linked to heavy metal stress induction. In the presence of heavy metals above a certain critical level, binding of metal to this metalloregulator releases it from its operon (de-repression) and triggers survival mechanisms to cope with this stress. A close examination of evolutionary analysis on SmtB/ArsR family members suggests a unifying overall fold and distinct metal selectivity profiles of one or two conserved metal coordination site(s). These two metal sites are designated α3 (i) and α5 (ii), named by the location of the metal binding sites within the known secondary structure of family members [41] (Fig. 2). For instance, the *α*3 metal binding site (i) has a highly conserved ELCV(C/G)D motif identified in members of the ArsR/SmtB family (Red box in Fig. 2) whereas the second metal binding site (ii) positioned between two protomers in dimer; Asp78, His79 of one protomer and His94 and Glu97 on the other protomer in *α*5 (cyan box in Fig. 2). However, the sequence analysis reveals that all antitoxins (Mab3862, Msmeg6762 and Rv2034) possess neither of the canonical metal-binding. For instance, Mab3862 has serine in the place of cysteine in ELCV(C/G)D consensus (*α*3 site). The mutation studies of its homologue, SmtB from *Synechococcus elongatus*, revealed that point mutation from cysteine to serine in *α*3 site might result in the loss of metal binding site but it still retain the ability to bind to DNA as regulator [42]. Therefore, the absence of the canonical metal box in antitoxin might have led to the complete loss of metal sensitivity, but it still functions as a transcription regulator by autoregulating of its operon. The phylogenic tree based on multiple sequence alignments of this family (Fig. 4) suggests that the most ancient ancestor is likely to have been one with the *α*3 binding site, before functional divergence and degeneration of the metal binding site resulted in the antitoxin group, whereas there is a re-emergence of the metal binding site in *α*5 on the other branch of the family [43].

**Fig. 2.**
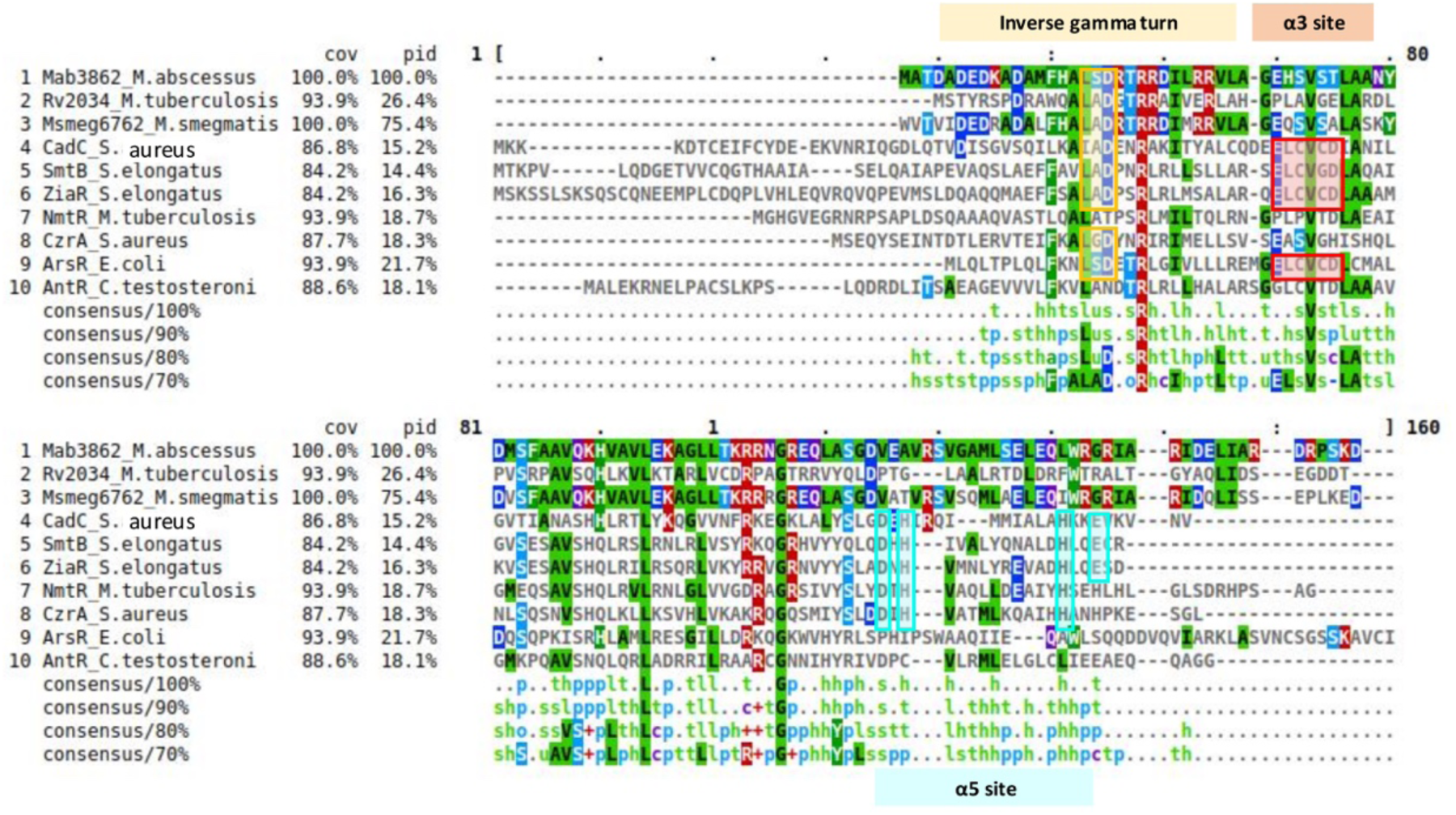
A multiple sequence alignment of several characterised ArsR/SmtB family members. GenBank accession in parenthesis: Mab3862 (ALM18125), Rv2034 (NP_216550), Msmeg6762 (ABK75527), CadC (P20047), SmtB (P30340), ZiaR (Q55940), NmtR (NP_218261), CzrA (O85142), ArsR (P15905) and AntR (WP034375793.1). Canonical metal binding sites on α3 (ELCV(C/G)D) and α5 (D,H on one protomer and H, E on the other protomer) are highlighted in red and cyan, respectively. The tripeptide LXD of the inverse gamma turn is noted in a yellow box.

Another structural difference between Mab3862 and other classical ArsR/SmtB family members is the absence of beta strands between *α*4 and *α*5. This region in many homologues of this family forms anti-parallel beta strands (Fig. 1d). It might be due to highly flexible regions (high B-factors) unable to assign any secondary structure in this crystal conditions but repeated data collection around this resolution (2.7Å) does not manifest any well-defined beta strands.

### 3.3 DNA binding site and domain prediction

Expression of TA system is tightly regulated by antitoxin by binding to inverted repeats located in the operon to repress its transcription [44]. Around the region between -10 and -35 upstream to the coding gene display nine inverted base pairs of hairpin DNA where Pribnow box overlaps (Fig. 3a). This hypothetical binding site might suggests that the Mab3862 repression is mediated by occlusion of promoter region by binding to this region. From consideration of other winged-helix protein-DNA complex structures, the putative DNA binding domain of this winged-helix protein consists of a wing region between two helices (*α*4 and *α*5) (green in Fig. 3b) and a face of *α*4 in Mab3862 (beige in Fig. 3b). There are two major DNA binding mechanisms exhibited by wHTH superfamily and they are exemplified by HNF-3 (hepatocyte nuclear factor 3) and RFX1 (regulatory factor X-box 1) respectively [45]. The overall topology and charge distribution (Fig. 3b) suggest that mab3862 might adopt the DNA binding mode similar to that of RFX1. The RFX1 shows unusual DNA binding mechanisms of wHTH superfamily; sequence-specific recognition is mediated mostly by interaction between major groove of DNA and wing region which mostly consists of basic residues (magenta in Fig. 3c) while a face of alpha helix (purple in Fig. 3c) shows largely neutral surface interacts with minor groove in the RFX1-DNA structure. Furthermore, biochemical studies and structural information of RFX1-DNA complex proves they bind to the double helical DNA with 2 fold crystallographic symmetry but there are little to no intermolecular interactions between monomers [46]. In the absence of direct protein-protein interactions, cooperativity appears to be mediated by protein-induced DNA deformation [46]. Thus, docking of Mab3862 to its promoter region with a guidance of its homologs indicates symmetrical binding of two Mab3862 to DNA cooperatively by conformational change in which wing gets nudged into major groove (red arrows in Fig. 3c) and helical structure of B-DNA gets slight distortion.

**Fig. 3.**
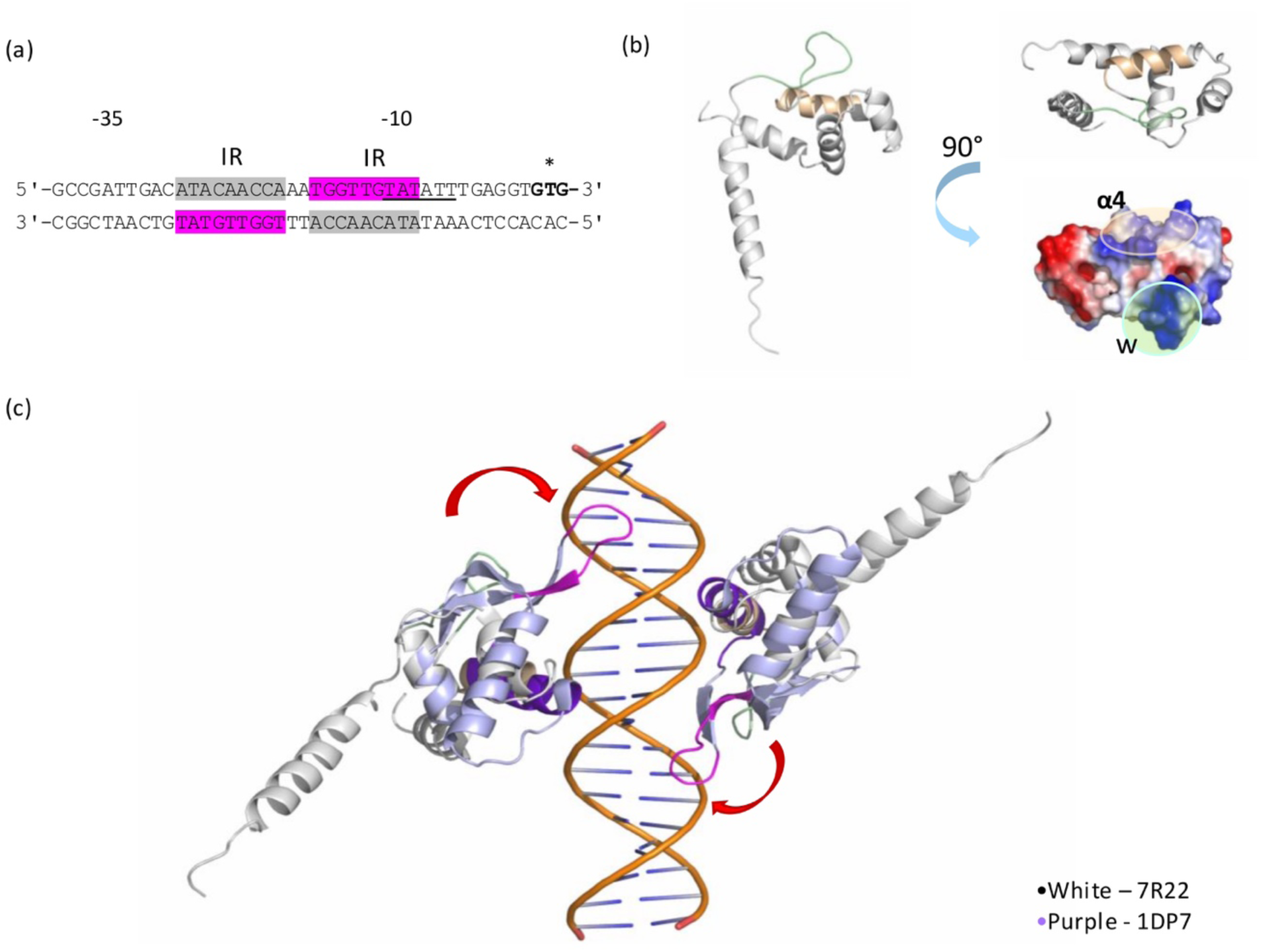
Mab3862 acts as an autoregulator by repressing its promoter region. a) putative palindromic sequence consists of nine base pairs and two base gaps between inverted repeats (IR) located between -30 and -10. A Pribnow box (TATATT) around -10 is noted by underline and a start codon (GTG) of Mab3862 gene is marked with asterisk. b) DNA binding domain of negative, blue denotes positive and white denotes neutral electrostatic potential regions consecutively. c) putative DNA fragments with sequence shown in (a) were generated computationally in B-DNA format. The docking of two Mab3862 were guided by the use of homolog (hRFX1, PDB id : 1DP7) DNA complex structure for correct docking conformation (ratio of DNA to protein 1:2). Red arrows predicts conformational change of wing region towards major groove upon DNA binding. Mab3862 coloured with green for wing region (w) between α4 and α5 and with orange for α4. Surface electrostatic properties of this protein also presented and labelled below; red denotes

**Fig. 4.**
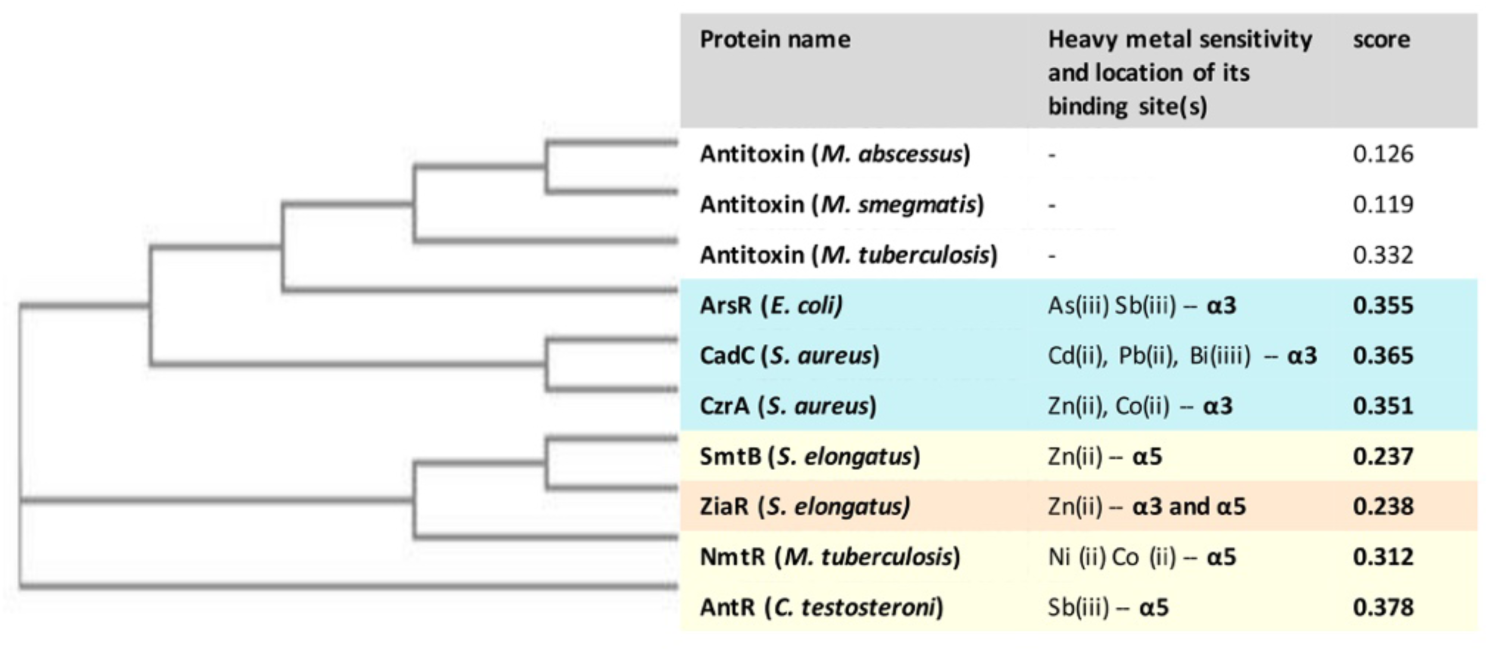
A phylogenetic tree of ArsR/SmtB family members based on the multiple sequence alignment shown in Fig. 2 along with metal sensitive profile and its binding location, created using the neighbour-joining method (NJ) and visualised with Clustal Omega and excel. The score represents evolutionary distances between protein members. Metal binding sites on α3 and α5 appeared to cluster separately on nodes of the dendrogram and are linked by a common evolutionary ancestor (except ZiaR, which has acquired both metal binding sites). The three antitoxins are more related to metalloregulator with α3 binding sites rather than α5 binding sites, suggesting the common ancestor might have had α3 binding sites that were then gradually lost in evolution.

### 3.4 Fragment screening using biophysical techniques

Having the structure of the target elucidated, we initiated a structure-guided FBDD effort targeting *M. abscessus* antitoxin binding sites by screening the in-house library of 480 small molecule fragments. Two biophysical methods, thermal shift assay (TSA) and plasmon resonance (SPR), were conducted in parallel to maximises the positive hits while minimising the false positives from double screenings (Fig. 5).

**Fig. 5.**
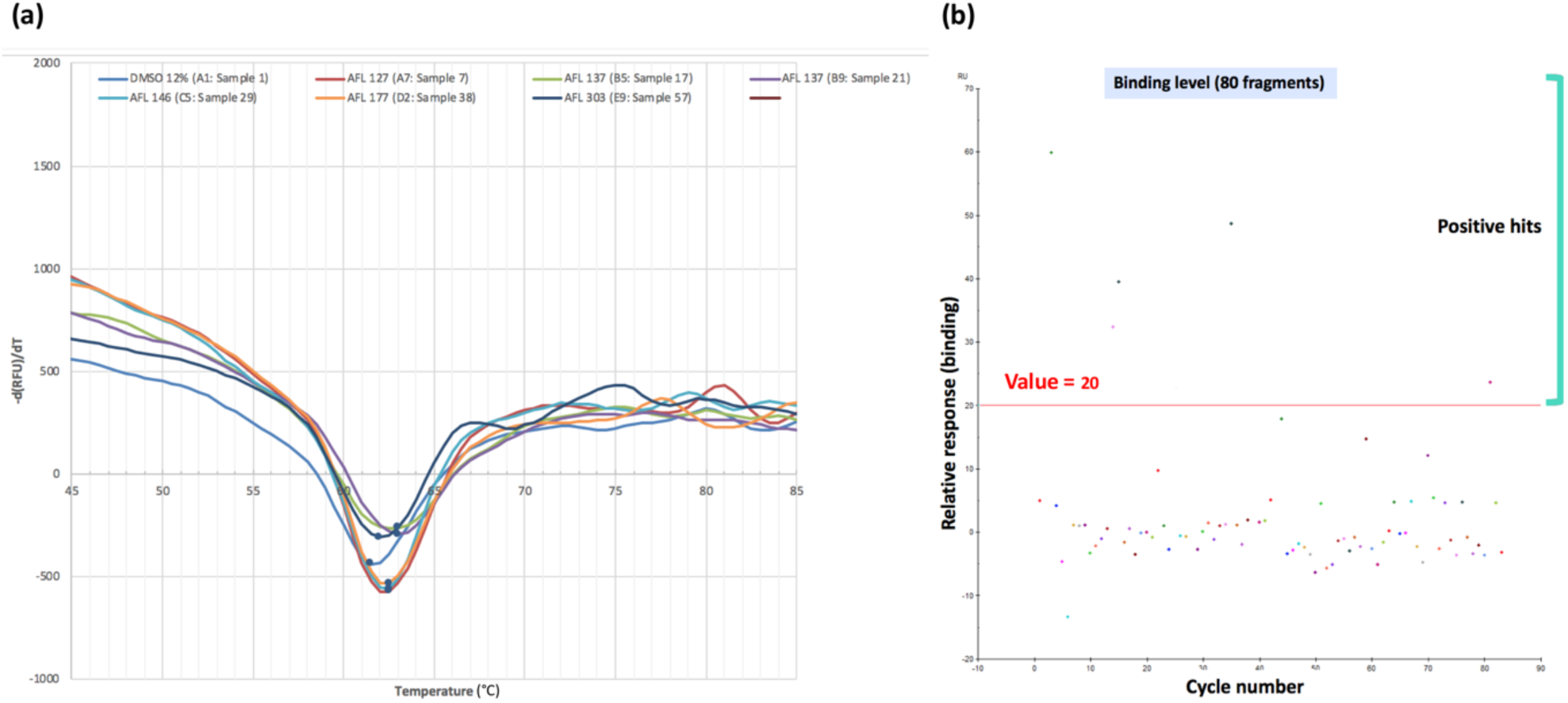
Examples of representative data from fragment-based screening techniques, (a) TSA and (b) SPR. (a) Fluorescence-based thermal shift has been used to measure the induced shift in the melting temperature of a protein in the presence of a fragment. Several fragments that showed a shift compared to the control (dark blue – only DMSO) were inspected in a graph. (b) Surface plasmon resonance (SPR) sensorgrams in which each point represents a different fragment being tested. Only those that showed responses significantly above the threshold were pooled. Here, the cut-off value used was 20 after solvent correction and subtracting the blank signals.

#### 3.4.1 TSA

Fluorescence-based thermal shift (TSA) detects molecules that increase the unfolding temperature of a target protein (ΔTm) by binding to and stabilizing the folded state [14]. Therefore, a shift in the melting point of a protein-ligand complex indicates stability in relation to its apo state (negative control). The thermal unfolding process is typically monitored in a plate-based format via an exogenous environmentally sensitive fluorescent dye (Sypro Orange). Because fragment binding is weak, the expected shifts tend to be small therefore, it requires highly sensitive detection equipment. The hits from TSA were identified as within a thermal shift cut-off value of 3 standard deviations from the negative control and their shift range from 0.5-1.0°C from the averaged control value (Fig. 5a). From this screening, 5 out of 480 small molecule fragments from library were validated through multiple repeated experiments.

#### 3.4.2 SPR

For Surface Plasmon Resonance (SPR) fragment screening, the target biomolecule is covalently linked to the gold surface of an SPR biosensor chip via amine coupling and solutions of single fragments are sequentially passed over it (1mM to 3mM). The cut-off was determined manually based on the overall level of response gained by visual inspections of graphs (Fig. 5b). When fragments bind to the immobilized target, increase in the surface mass therefore refractive index is monitored in real time. From the time-dependent fragment association−dissociation response, the binding kinetics can be measured and the binding affinity calculated. From this screening, 26 out of 480 fragments were tested positive through duplicate experiments.

Surprisingly, there was no overlap of positive hits between TSA and SPR results. Previous studies on six different biophysical screening including SPR, TSA, NMR, biochemical assays and others on a single target revealed a disappointingly low overlap between methods with no single common hit [47]. Each technique operating on different mechanisms under various conditions (e.g. buffer, temperature, concentration of protein etc.) ends up measuring and detecting different set of binders. However, use of multiple methods in parallel is still encouraged to maximise hits which potentially could lead to a genuine hit confirmed by X-ray crystallography.

### 3.5 Fragment screening using *in silico* docking

Virtual screening can identify the binding mode for a biophysically validated hit in the absence of structural information about the protein-ligand complex. For molecular docking to be useful, it must be able to produce the correct binding mode and evaluate it based on binding energy. The empirical scoring function (Glide Score) computes approximate free energy of ligand and can be used to rank multiple poses of the same or different ligands in virtual screening [31]. Initially, the searching of potential binding site of Mab3862 has led to two sites where fragments are likely to bind with site score roughly similar to each other (0.92 and 0.90 for site I and site II, respectively) (Fig. 6). Out of 480 fragments, hits identified from TSA and SPR were compiled as one list (Table 3) and the mode of binding examined *in silico* carefully. Some of the examples that scored highly both in experiment and *in-silico* docking are presented in Fig. 7. Then, the common binding sites with overlapping interacting residues were grouped together to draw a common chemical scaffold to build a pharmacophore model (Fig. 8). A pharmacophore proposes necessary three-dimensional features for recognition of a ligand by a biological target. In this approach, energies derived from docking (Glide scores) are mapped onto all atoms that define six pharmacophore chemical features in the hypothetical binding pocket; hydrogen bond acceptor, hydrogen bond donor, hydrophobic, negative ionizable, positive ionizable, and aromatic ring features. The hydrogen bond donors and acceptors were assigned based on pure projected points or vectors, leading to different projected points-based and vector-based hypotheses. This can be used to manually design new compounds or to search a database to find molecules that contain this required features [48]. This will assist the medicinal chemist to work on rational design of drug synthesis from fragments with some structural insights.

**Fig. 6.**
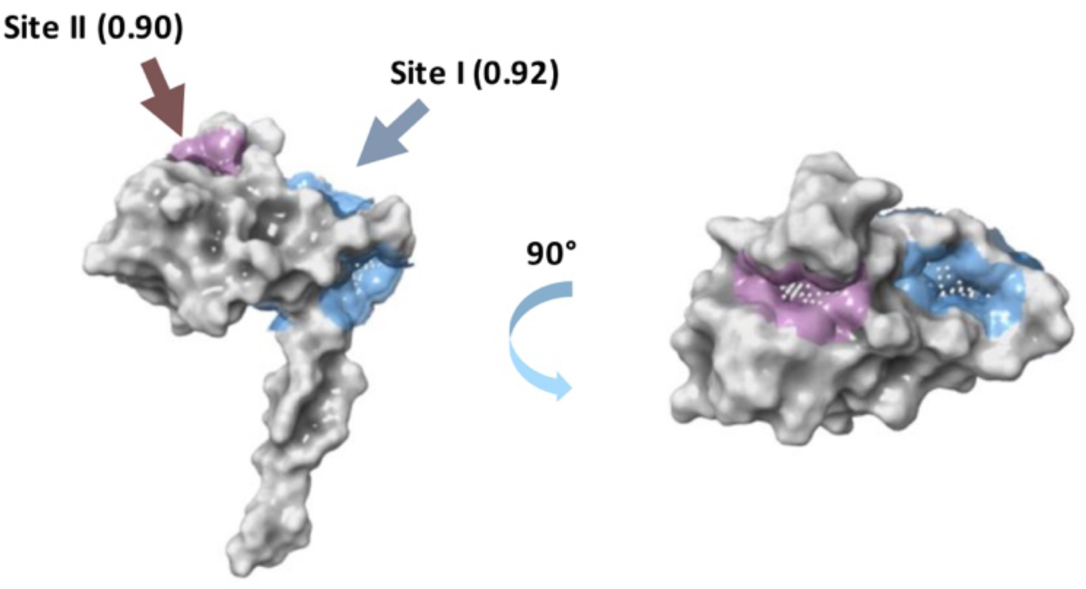
Site map of Mab3862. Two sites were identified (site I and site ii) with similar site scores annotated in brackets. A score above 1.0 means an excellent binding site whereas score between 0.8 and 1.0 indicates challenging but still valid binding site. A score below 0.8 means non-druggable

**Fig. 7.**
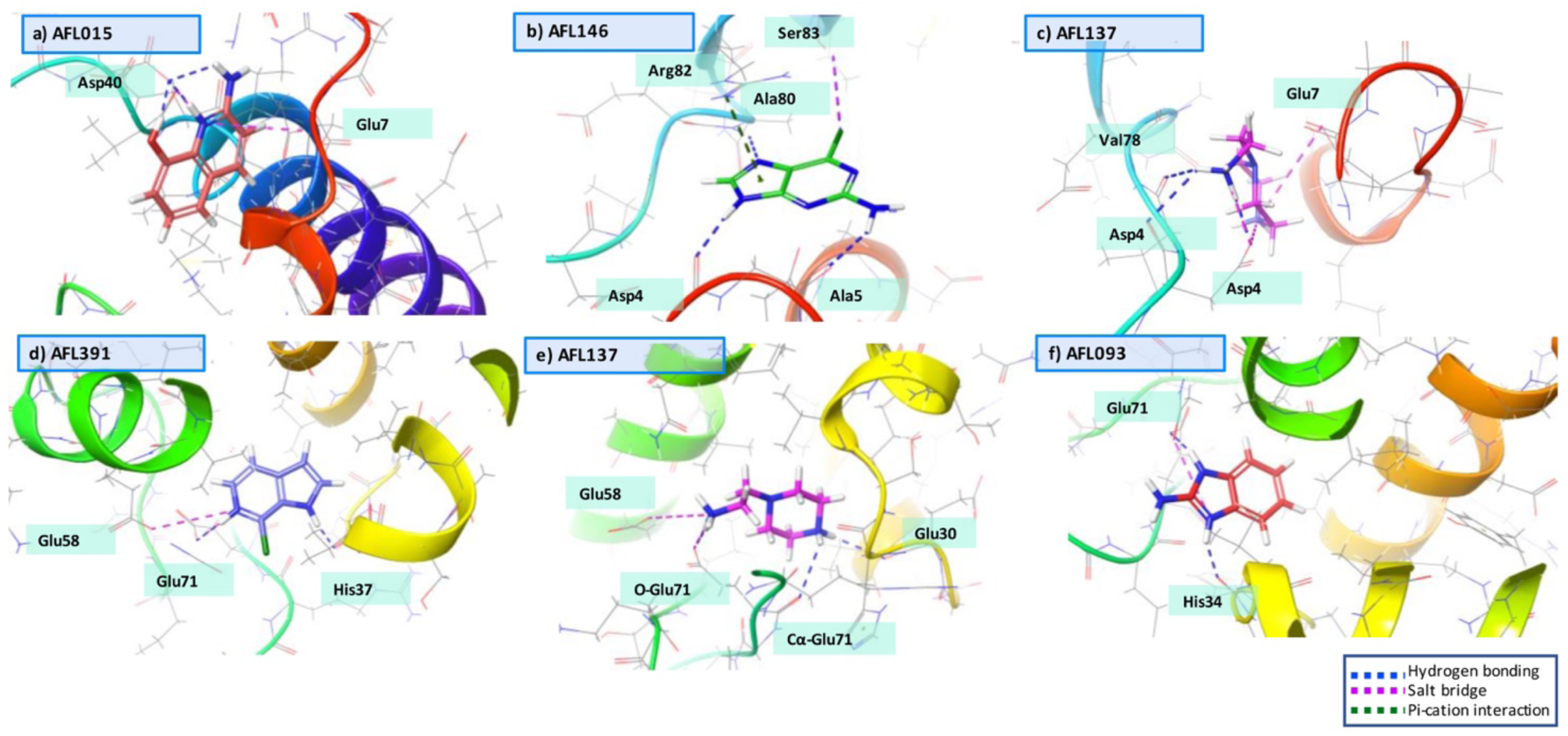
Examples of *in silico* molecular docking, which produces ranked and docked conformations of fragments from the in-house library at site 1 (A-C) and site2 (D-F). Each fragment was labelled on top left, along with residues interacting with fragments. Hydrogen bonding in blue, salt bridges in magenta and Pi-cation interactions in green are shown clearly with dash lines.

**Fig. 8.**
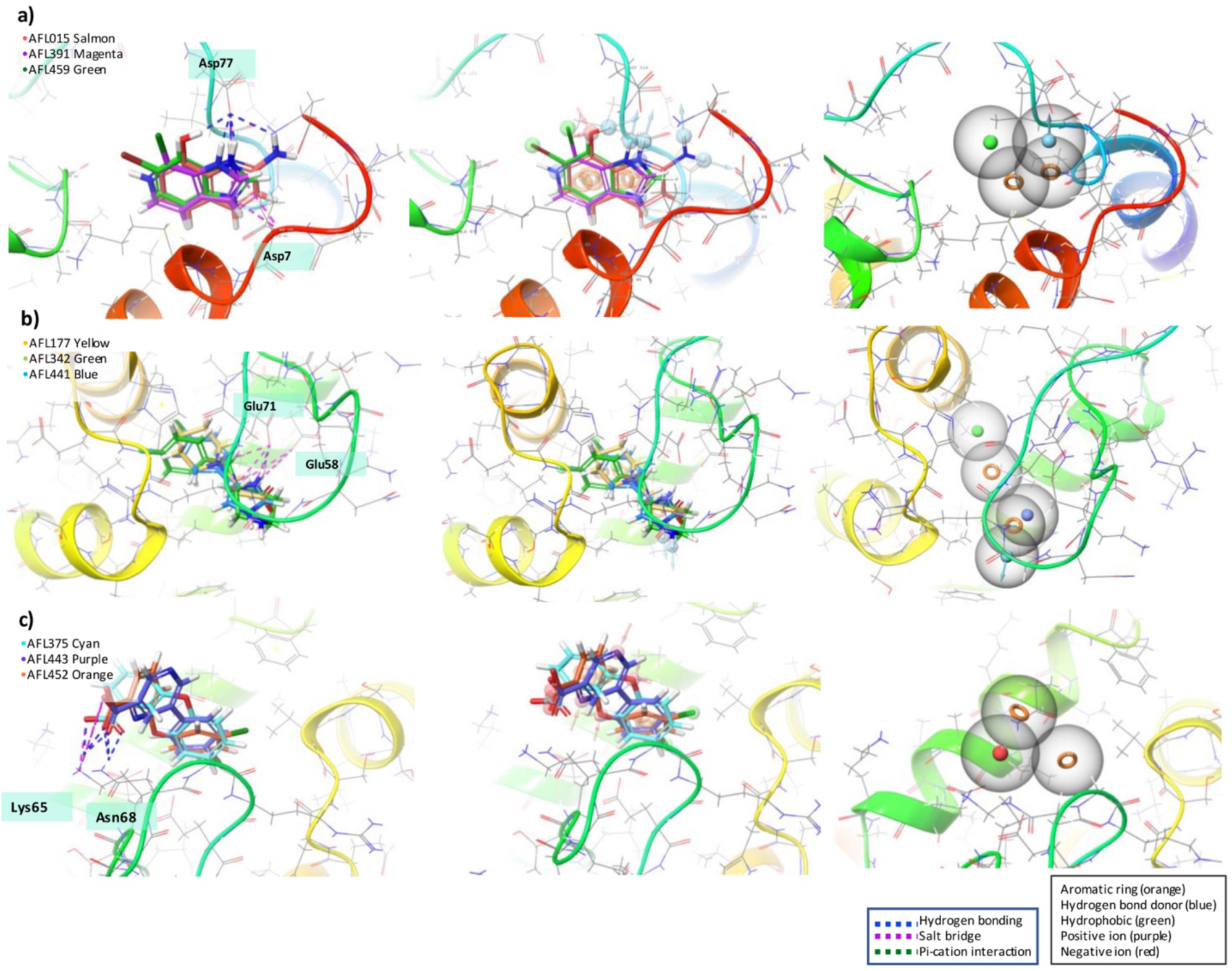
Fragments in groups of three were analysed based on the common residues which they interact at each site. The fragments were superimposed to create the pharmacophore models. An orange hollow ring represents an aromatic ring; light blue presents hydrogen bond donors; green is for a hydrophobic moiety, purple for a positive ion and red for negative ions. This Fig. was generated by Schrödinger software.

## 4. Conclusions

Comparative genomic studies between epidemic and non-epidemic species showed that the presence of TA modules has a significant association with the pathogenic repertoire [49]. For example, *M. tuberculosis* contains at least 30 functional TA genes [50], whereas its non-pathogenic counterpart, *M. smegmatis* has only three [51,52]. However, the drug discovery process of targeting TA is not as straightforward as classical approaches that tended to target active sites [12]. For instance, targeting protein-protein interactions (PPI) between toxin and antitoxin has always been challenging due to the small size of the protein and poorly defined intermolecular contact area. Also, the redundancy of multiple TA systems in bacteria makes it unclear whether inhibiting a single TA system (toxin) will be sufficient to either kill or prevent growth of bacteria; rather, it might require a cocktail of inhibitors [11].

Several reports discuss druggability of toxin-antitoxin complexes and introduce various methods of targeting TAs [11,12]. The ultimate aim of drug discovery is to release toxin from its partner either by preventing formation or disrupting the TA complex in order to proceed a toxic effect on the bacterial cell (Fig. 9). If the function of the toxin is known to be bactericidal (e.g. by blocking DNA replication, inhibiting cell wall synthesis), it would kill bacteria either on its own or as a group of toxins down the cascade pathway leading to cell death. On the other hand, if the function of toxin is bacteriostatic, for instance, by inducing dormant state of persistence, one might take advantage of this situation by employing PZA drugs [53]. Pyrazinamide (PZA), a nicotinamide analogue, is a unique drug that kills non-replicating persisters in many pathogenic species. It has been reported in *M. tuberculosis* infection that the use of PZA shortened the treatment period by 3 to 6 months by getting rid of persistent subpopulations that are not killed by any other drugs [53].

**Fig. 9.**
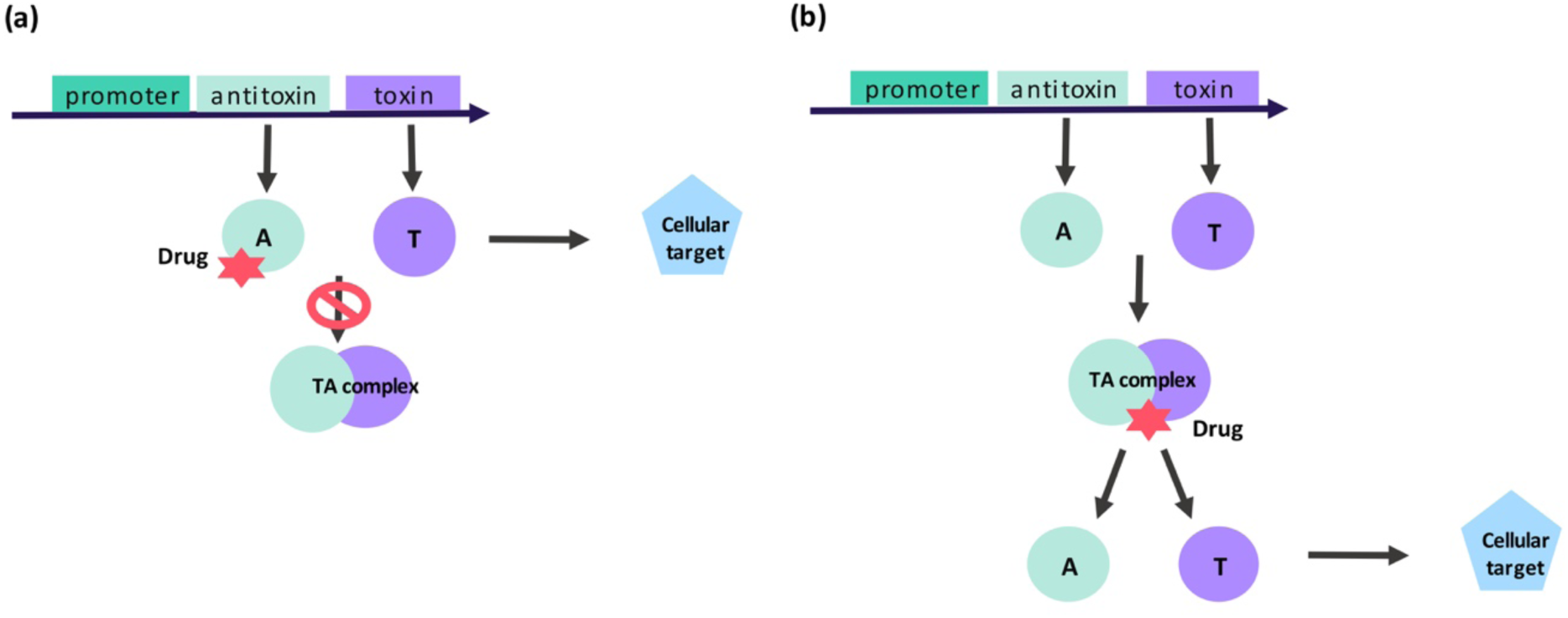
Different antibacterial strategies leading to toxin activation (a) prevention of TA complex formation and (b) disruption of TA complexes. Cellular target and its reaction depends on the type of toxin. They are known to be involved in post-segregational killing (PSK), abortive infection, biofilms and persister cell formation. A stands for antitoxin and T stands for toxin.

In conclusion, the main focus of targeting TA should remain on separating toxin from its cognate antitoxin by either blocking or disrupting the complex. By using a fragment-based drug discovery with *in silico* guidance, we have discovered a few valid hits that bind to novel sites on *de novo* structure of Mab3862 antitoxin. The data presented here clearly demonstrate the potentiality and scope to target of antitoxin from TA complex with dedicated drug discovery programs and we propose toxin and antitoxin as the optimal candidates for fighting with persisters and growing resistance. Indeed it is noteworthy that pivoting the effort towards exploring unconventional and novel targets would broaden the spectrum of classes of antibiotics, therefore diversifying the treatment and therapy depending on types of infection.

## Abbreviations

Mab: *Mycobacterium abscessus*
NTM: non tuberculous mycobacteria
TA: Toxin and antitoxin
PPI: protein-protein interaction
FBDD: fragment-based drug discovery
CF: cystic fibrosis.

## Acknowledgement

This work is funded by the Cystic Fibrosis Trust (Registered as a charity in England and Wales) and National Research Foundation of Korea (NRF) through a grant funded by the Korean Government (MEST) [2018R1A5A2024425, 2021R1F1A1050961].

The authors thank the Diamond Light Source for beam-time and the staff of beamlines I04 for assistance with data collection. In-house fragment library was kindly offered by Chris Abell’s group in a form of a long-term collaboration between the Blundell and Abell groups.

