## Supplementary material for "Structural-guided fragment-based drug discovery applied to antitoxin, MAB3862 opens a new possibility of exploring the Toxin and Antitoxin for antibiotics": TABLES

**Table 1. Statistics of data collection and refinement of Mab3862 (7r22)**

| **Data collection and processing** | |
|---|---|
| Diffraction source | I04 beamline at Diamond Light Source |
| Wavelength | 0.9795 Å |
| Temperature | 100K |
| Detector | PIXEL (DECTRIS EIGER2 XE 16M) |
| Total rotation range per image | 0.1° |
| Total rotation range | 360° |
| Exposure time per image | 0.008s |
| Space group | $P4_132$ |
| Unit cell parameters | 106.89 Å 106.89 Å 106.89 Å<br><br>90.00° 90.00° 90.00° |
| Resolution range | 47.80-2.73 Å |
| Rmerge | 0.12 |
| Total reflections | 405313 |
| Unique reflections | 5963 |
| Multiplicity | 68 |
| <I/σ> | 0.6 at 2.73Å |
| Overall B factor from Wilson plot | 94.0 $Å^2$ |
| **Structure refinement** | |
| Resolution range | 2.72-2.76 Å |
| Completeness in resolution range | 75.5% |
| Working set reflections | 4059 (90%) |
| Test set reflections | 451 (10.02%) |
| Final $R_{work}$ | 0.347 |
| Final $R_{free}$ | 0.299 |
| Molecules in asymmetric unit | 1 |
| RMSD Bond/Angles | 0.6013°/ 2.036 Å |
| **Ramachandran plot** | |
| Favoured regions | 68% |
| Allowed | 17% |
| Outliers | 14% |
| Rotamer outliers | 6% |
| Solvent contents | 72.46% |
| Matthew coefficient | 4.47$Å^3$ |
| Average B factor | 98.3 $Å^2$ |

**Table 2. Mab3862 dimer interface interaction**

| | | |
|---|---|---|
| **Number of residues** | | |
| Interface | 36 | 35.6% |
| Surface | 100 | 99.0% |
| Total | 101 | 100.0% |
| **Solvent accessible area (Å)** | | |
| Interface | 1454.1 | 19% |
| Total | 7633.4 | 100% |
| **Solvation energy, kcal/mol** | | |
| Isolated structure | -62.8 | 100% |
| Gain on complex formation | -13.5 | 21.5% |
| Average gain | -3.4 | 5.5% |
| P-value | 0.017 | - |

**Table 3. A list of putative positive hits from SPR and TSA with 2d structure and glide score annotated in each site.**

| AFL Sample | Source | site 1 | site 2 | chemical name | structure 2D |
|---|---|---|---|---|---|
| 15 | SPR | -7.073 | -4.979 | 2-Amino-8-quinolinol | 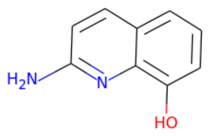 |
| 447 | SPR | -6.966 | -5.047 | 5-Bromo-2-(hydroxymethyl)indole | 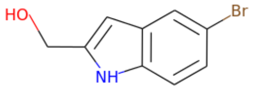 |
| 137 | TSA | -6.496 | -6.177 | 1-(2-Aminoethyl)piperazine | 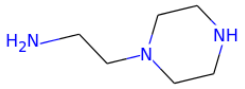 |
| 93 | SPR | -6.298 | -5.824 | 3,4-Dichlorobenzylamine | 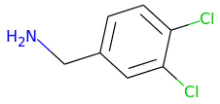 |
| 459 | SPR | -6.293 | -5.474 | 6-Bromo-1H-benzimidazole | 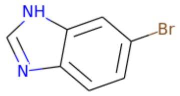 |

| | | | | | |
|---|---|---|---|---|---|
| 428 | SPR | -6.235 | -5.198 | 4-Phenylbenzylamine, 97% | 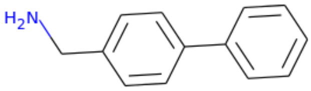 |
| 391 | SPR | -6.177 | -7.412 | 7-Chloro-6-azaindole | 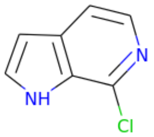 |
| 277 | SPR | -5.920 | -3.936 | 4-Aminoquinaldine | 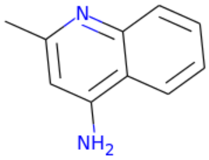 |
| 146 | TSA | -5.889 | -4.923 | 2-Amino-6-chloropurine | 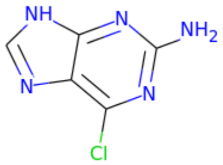 |
| 441 | SPR | -5.525 | -5.185 | 4-(4-Chlorophenyl)-1H-imidazole | 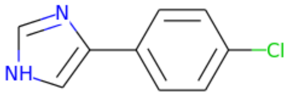 |

| | | | | | |
|---|---|---|---|---|---|
| 201 | SPR | -5.383 | -4.292 | 5-Methoxy-3-indolylacetonitrile | 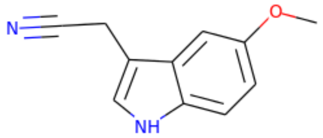 |
| 303 | TSA | -5.329 | -4.727 | 1-Acetyl-4-(4-hydroxyphenyl)piperazine | 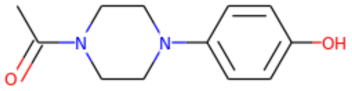 |
| 65 | SPR | -5.221 | -5.435 | 3-(4-Chlorophenyl)-1H-pyrazol-5-amine | 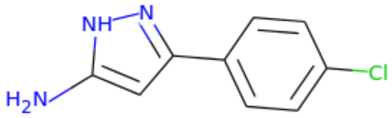 |
| 158 | SPR | -5.182 | -4.692 | 4-Amino-1-benzylpiperidine | 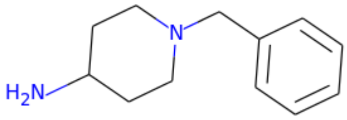 |

| | | | | | |
|---|---|---|---|---|---|
| 175 | SPR | -5.137 | -4.430 | N-Methyl-N-(2-thien-2-ylbenzyl)amine | 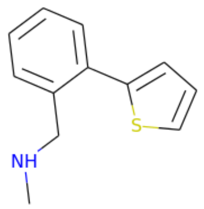 |
| 127 | TSA | -5.096 | -5.408 | 4-Piperidinemethanol | 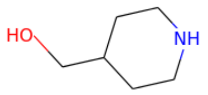 |
| 16 | SPR | -5.092 | -5.995 | 2-Aminobenzimidazole | 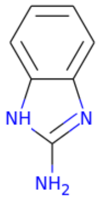 |
| 80 | SPR | -4.966 | -4.986 | 6-Hydroxy-2,3-dihydrobenzo[b]furan-3-one | 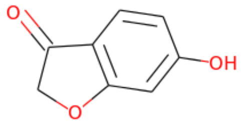 |
| 352 | SPR | -4.948 | -4.290 | Indole-3-carbonitrile | 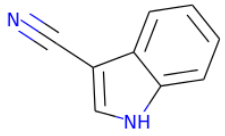 |

| | | | | | |
|---|---|---|---|---|---|
| 177 | TSA | -4.922 | -4.694 | 1-Piperidinepropionic acid | 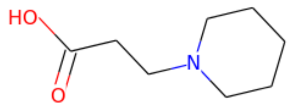 |
| 452 | SPR | -4.809 | -4.000 | 6-(3-Chlorophenoxy)pyridine-2-carboxylic acid | 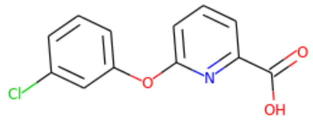 |
| 342 | SPR | -4.795 | -4.394 | 1-(4-Fluorobenzyl)homopiperazine | 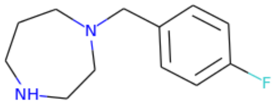 |
| 373 | SPR | -4.779 | -3.740 | 5-Chlorobenzoxazole | 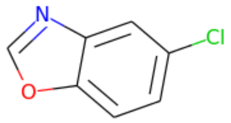 |
| 66 | SPR | -4.770 | -4.647 | 3-(Phenoxymethyl)benzoic acid | 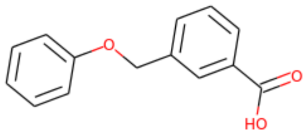 |

| | | | | | |
|---|---|---|---|---|---|
| 443 | SPR | -4.761 | -3.959 | 5-(4-Chlorophenyl)pyridine-3-carboxylic acid | 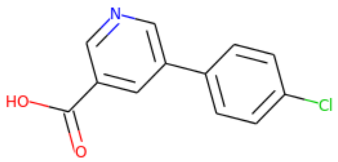  |
| 245 | SPR | -4.641 | -5.007 | 3-(4-Methylphenyl)-1H-pyrazol-5-amine | 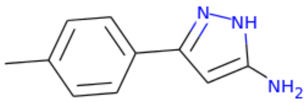  |
| 450 | SPR | -4.537 | -4.105 | 5-Methoxy-1H-pyrrolo[2,3-c]pyridine-2-carboxylic acid | 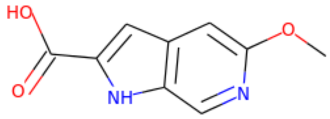  |
| 449 | SPR | -4.192 | -3.957 | 4-(4-Trifluoromethoxyphenyl)picolinic acid | 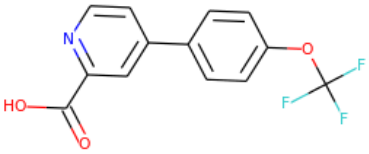  |

| | | | | | |
|---|---|---|---|---|---|
| 375 | SPR | -4.174 | -4.016 | 3-Phenoxybenzoic acid | 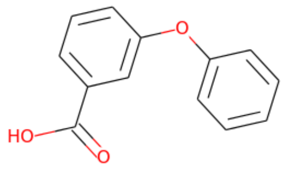 |
| 448 | SPR | -3.986 | -4.058 | 5-Chloroindole-2-carboxylic acid | 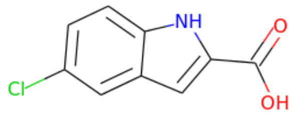 |
| 387 | SPR | -3.553 | -4.549 | 4-Phenoxyphenylacetic acid | 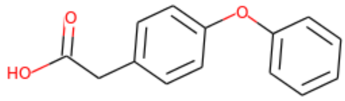 |
